# Population Genetic Analysis of the DARC Locus (Duffy) Reveals Adaptation from Standing Variation Associated with Malaria Resistance in Humans

**DOI:** 10.1101/050096

**Authors:** Kimberly F. McManus, Angela Taravella, Brenna Henn, Carlos D. Bustamante, Martin Sikora, Omar E. Cornejo

## Abstract

The human DARC (Duffy antigen receptor for chemokines) gene encodes a membrane-bound chemokine receptor crucial for the infection of red blood cells by *Plasmodium vivax*, a major causative agent of malaria. Of the three major allelic classes segregating in human populations, the FY*O allele has been shown to protect against *P. vivax* infection and is near fixation in sub-Saharan Africa, while FY*B and FY*A are common in Europe and Asia, respectively. Due to the combination of its strong geographic differentiation and association with malaria resistance, DARC is considered a canonical example of a locus under positive selection in humans.

Here, we use sequencing data from over 1,000 individuals in twenty-one human populations, as well as ancient human and great ape genomes, to analyze the fine scale population structure of DARC. We estimate the time to most recent common ancestor (T_MRCA_) of the FY*O mutation to be 42 kya (95% CI: 34–49 kya). We infer the FY*O null mutation swept to fixation in Africa from standing variation with very low initial frequency (0.1%) and a selection coefficient of 0.043 (95% CI:0.011–0.18), which is among the strongest estimated in the genome. We estimate the T_MRCA_ of the FY*A mutation to be 57 kya (95% CI: 48–65 kya) and infer that, prior to the sweep of FY*O, all three alleles were segregating in Africa, as highly diverged populations from Asia and ≠Khomani San hunter-gatherers share the same FY*A haplotypes. We test multiple models of admixture that may account for this observation and reject recent Asian or European admixture as the cause.

**Author Summary:** Infectious diseases have undoubtedly played an important role in ancient and modern human history. Yet, there are relatively few regions of the genome involved in resistance to pathogens that have shown a strong selection signal. We revisit the evolutionary history of a gene associated with resistance to the most common malaria-causing parasite, *Plasmodium vivax*, and show that it is one of regions of the human genome that has been under strongest selective pressure in our evolutionary history (selection coefficient: 5%). Our results are consistent with a complex evolutionary history of the locus involving selection on a mutation that was at a very low frequency in the ancestral African population (standing variation) and a large differentiation between European, Asian and African populations.

## Introduction

Infectious diseases have played a crucial part in shaping current and past human demography and genetics. Among all infectious diseases affecting humans, malaria has long been recognized as one of the strongest selective pressures in recent human history [1,2]. The Duffy antigen, also known as DARC (Duffy antigen receptor for chemokines) and more recently as ACKR1 (atypical chemokine receptor 1), is a transmembrane receptor used by *Plasmodium vivax*, a malaria-causing protozoan, to infect red blood cells. *P. vivax* causes a chronic form of malaria and is the most widespread type of malaria outside of Africa [3,4].

The *DARC* gene has three major allelic types that are the product of two common polymorphisms, forming the basis of the Duffy blood group system [5,6]. The two variant forms, FY*B and FY*A, are the allelic types commonly observed in non-African populations. FY*B is the ancestral form of the receptor, and is widespread in Europe and parts of Asia. FY*A is defined by a derived non-synonymous mutation (D42G, rs12075) in the *P. vivax* binding region of the DARC protein. It is the most prevalent of the three alleles in modern human populations, with highest frequency in Asia (predicted frequency >80%) and at 30-50% frequency in Europe [4]. FY*A is also present in southern Africa, despite absence from western and central Africa [4,7–9]. FY*O (also known as Duffy null) is defined by a mutation (T-42C, rs2814778) in the GATA-1 transcription factor binding site in the DARC gene promoter region, and occurs mostly on a FY*B background. The derived FY*O mutation exhibits extreme geographic differentiation, being near fixation in equatorial Africa, but nearly absent from Asia and Europe [4].

Of the three allelic types, FY*A and FY*B are functional proteins, while FY*O does not express the protein on erythrocyte surfaces due to a mutation in the promoter region, which causes erythroid-specific suppression of gene expression [6,10]. The lack of expression of DARC in erythrocytes has been shown to halt *P. vivax* infection [6,10]. Moreover, recent evidence shows that heterozygous individuals have reduced DARC gene expression and evidence of partial protection against *P. vivax* [11,12]. It has been proposed that due to the near-fixation of FY*O, *P. vivax* infection in humans is largely absent from equatorial Africa. Phenotypic implications of the FY*A mutation are less clear than FY*O; however, there is evidence of natural selection and reduced *P. vivax* infection in individuals with this genotype ([13,14], conflicted by reports in the Brazilian Amazon however [12,15,16]). An important recent discovery suggests low levels of *P.vivax* infection in FY*O homozygotes [17–21], which indicates that *P. vivax* might be evolving escape variants able to overcome the protective effect of FY*O.

There is long running interest characterizing the evolutionary forces that have shaped the Duffy locus. The combination of strong geographic differentiation and a plausible phenotypic association (resistance to malaria) has led to the Duffy antigen being cited as a canonical example of positive selection in the human genome (eg. [22–24]); however, details of its genetic structure remain understudied. Though touted as under positive selection, the few early population genetic studies of this locus found complex signatures of natural selection [25,26] and it is rarely identified in whole genome selection scans [27–35]. Some genomic loci display signatures of selection readily captured by standard methods, yet other well-known loci, like FY*O, are overlooked potentially due to intricacies not captured by simple models of hard selective sweeps. Detailed analyses of the haplotype structure of these loci can help us better understand complicated scenarios shaping genetic variation in loci under selection.

What makes the evolution of FY*O such a complex and uncommon scenario? *Plasmodium* species and mammals have coexisted for millions of years, with frequent cases of host-shifts and host range expansions along their evolution [36,37]. Great apes are commonly infected with malaria-related parasites [38,39] and recent evidence suggests that human *P. vivax* originated in African great apes [38]; contrasting with previous results that supported an Asian origin for *P. vivax* [40,41]. In addition to the complex evolutionary relationship among *Plasmodium* species and mammals, the specific mechanisms of invasion of erythrocytes employed by different species are highly diverse and present commonalities among species. DARC erythroid expression also influences infection in a variety of other species. It is required for infection by *Plasmodium knowlesi*, a malaria parasite that infects macaques, and SNPs upstream of the *DARC* gene homologue in baboons influence DARC expression and correlate with infection rates of a malaria-like parasite [42,43]. Other studies utilizing single primate sequences found evidence for accelerated evolution in this gene region [44,45]. However, no previous study analyzed population level sequence data in great apes.

Despite the general understanding of the relevance of DARC in the evolution of the interaction between *Plasmodium* and primates, a thorough analysis of the complex evolutionary history of this locus using recently available large-scale genomic datasets of diverse human populations is still lacking. Here, we analyze the fine scale population structure of DARC using sequencing data from twenty-one human populations (eleven African populations), as well as great ape and ancient human genomes. We estimate the time to most recent common ancestor of the FY*A and FY*O mutations and estimate the strength of selection of FY*O. We propose a model of FY*O’s spread through Africa, which builds on previous findings and provides a more complete picture for the evolution of FY*O. We further explore the relationship between the common FY*A haplotype in Asia and the FY*A haplotype found in southern Africa. Lastly, we investigate selection and SNPs in great ape sequence data and provide the first population genetic analysis of DARC in apes.

## Results

### Population Genetics of the Duffy Locus

#### Geographical distribution

We observe broad consistency between the geographic distribution of the major allelic types in our dataset and previously published results [4] (Fig 1, S2 Table). We find that the FY*O mutation is at or near fixation in western and central African populations, but almost absent from European and Asian samples. All sampled sub-Saharan African populations show frequencies of >99% for FY*O, with the exception of the southern African Zulu and ≠Khomani San populations that contain all three of the FY*A, FY*B and FY*O alleles. FY*A is the dominant allele in all five Asian population samples (89-95%), while FY*B is most common in all five European populations (55-70%).

**Fig 1.**
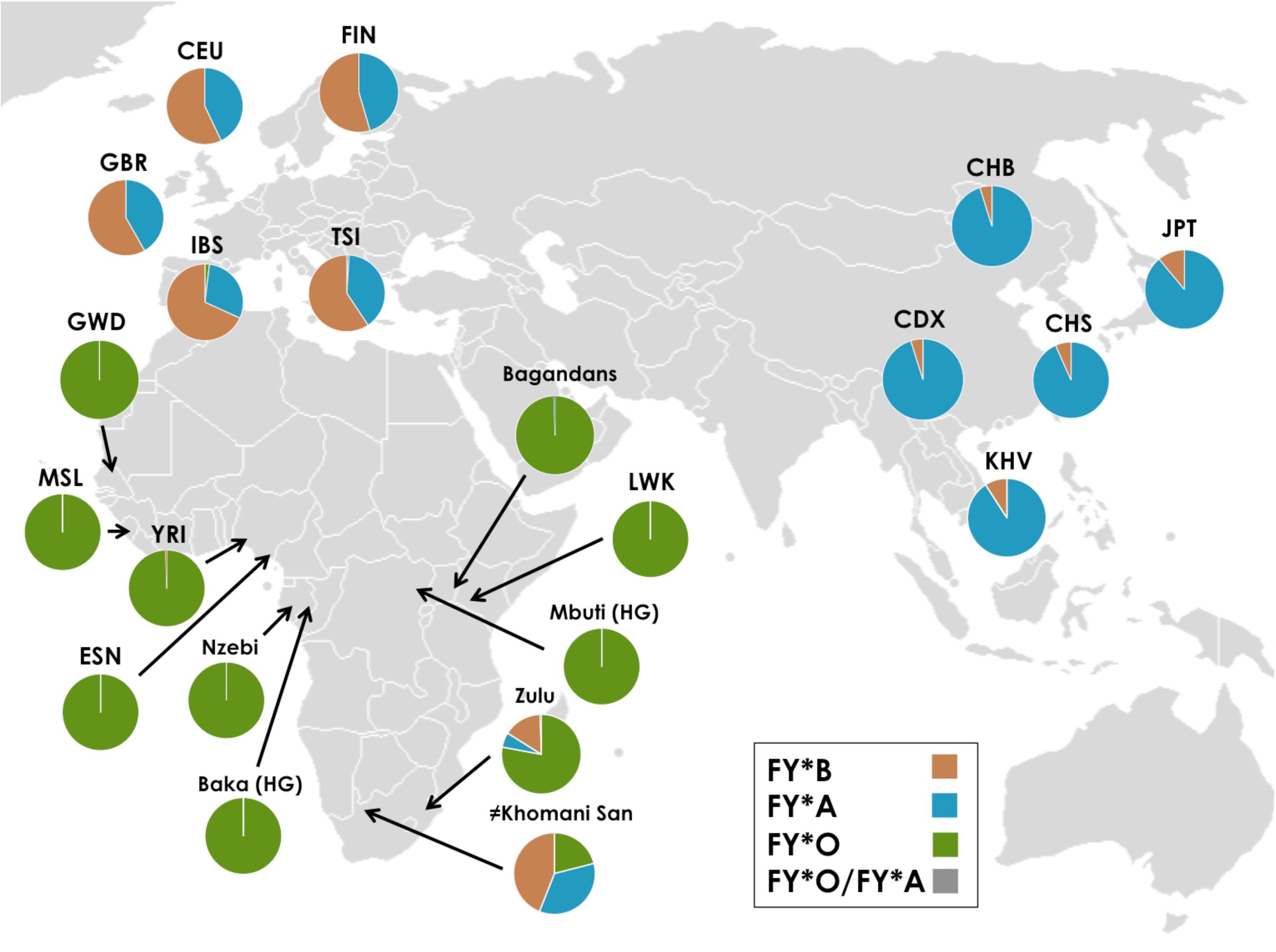
Geographical distribution of allelic classes in samples.

#### Phylogenetics

We surveyed the 5 kb region surrounding the FY*O mutation. The FY*A mutation is located 671 basepairs downstream of the FY*O mutation. Median-joining haplotype networks of this locus reveal decreased diversity in FY*O and FY*A haplotypes and little geographic structure within continents (Fig 2). We analyzed all unique haplotypes observed at least four times in this 5kb region and find FY*O and FY*A allelic classes form distinct clusters, while FY*B is more diverse. Recombination is observed on all haplotypes in this region.

**Fig 2.**
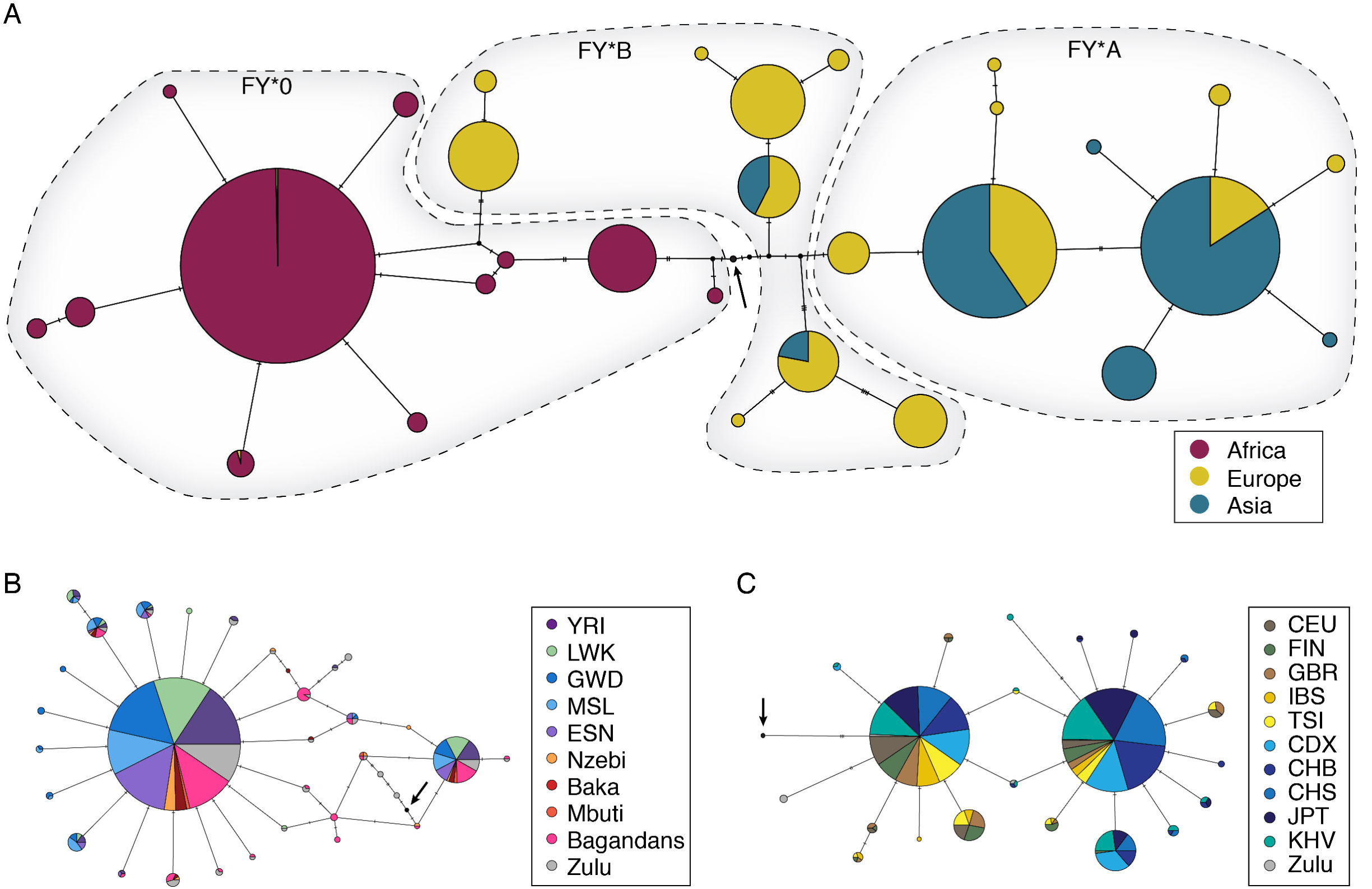
Haplotype Networks. Median joining networks of three subsets of haplotypes in the 5kb region centered on the FY*O mutation. The FY*A mutation is located 671 bps downstream from the FY*O mutation. Arrows indicates ancestral sequence. A) All haplotypes observed at least four times B) All FY*O haplotypes observed at least twice. (Note that none of the FY*O haplotypes in this network also carry the FY*A mutation.) C) All FY*A haplotypes observed at least twice. (Note that none of the FY*A hapotypes in this network also include the FY*O mutation.)

FY*O exhibits two major haplotypes, as seen previously [25], which are defined by four SNPs (chr1:159174095, chr1:159174885, chr1:159176831, chr1:159176856). The haplotypes are at unequal frequency with the most common haplotype at 86% frequency in FY*O sub-Saharan African samples, while the minor haplotype is at 10% frequency. FY*O’s haplotypes exhibit little to no population structure between African populations, though the most common haplotype is at slightly lower frequency in eastern Africa, compared with western and southern Africa. We also note there is evidence of bias between the two methods of SNPs calling (see Discussion). Notably, the FY*O haplotypes observed in the Baka and Mbuti hunter-gatherer populations are identical to Bantu African haplotypes, in stark contrast to the deep divergence between these populations at the genome-wide level [46].

The FY*A allele also exhibits two major haplotypes and reduced diversity relative to the ancestral FY*B allele. FY*A’s two common haplotypes are at more similar frequencies, though the more derived haplotype is dominant in Asia while the more ancestral haplotype is more common in Europe. There is significant recombination between FY*A and FY*B as, unlike FY*O, they coexist in many populations.

### Evidence of selection in DARC

#### Evidence of positive selection at FY*O

Despite FY*O’s biological support for positive selection, it has not been identified as a potential selected region in many genome-wide selection scans [27–35]. Accordingly, we find the DARC promoter region is not an outlier in the genome with respect to segregating sites, average number of pairwise differences nor Tajima’s D (S3 Table). Though the DARC promoter region has the fewest SNPs in African populations, it has more pairwise differences likely due to two divergent FY*O haplotypes in these populations.

To further investigate signatures of selection in the DARC region, we analyzed statistics from three main classes of selection scans: population differentiation (F_*ST*_), site frequency spectrum (Sweepfinder [47,48]), and linkage disequilibrium (H-scan [49]) (Table 1, S4-S6 Tables).

**Table 1.** Selection scan results

|  | 20KB |  | 100KB |  |
| --- | --- | --- | --- | --- |
|  | Sweepfinder | H-scan | Sweepfinder | H-scan |
| African | 0.052 | 0.076 | 0.032 | 0.054 |
| Asian | 0.303 | 0.613 | 0.155 | 0.494 |
| European | 0.582 | 0.701 | 0.626 | 0.252 |

Selection scan results for region around the FY*O mutation. Numbers indicate empirical p-value for Sweepfinder and H-scan statistics in 20kb and 100kb regions centered on the FY*O mutation. These are calculated by comparing the FY*O region statistics with the distribution of statistics from the whole genome. Table includes an average of results from each of the fifteen 1000 Genomes populations and results compare regions of similar recombination rate.

We find that FY*O has the largest population differentiation, as measured by F_*ST*_, of any SNP in the genome among the 1000 Genomes populations. This signature extends to the 100 kb region surrounding FY*O, though it is reduced to the 96.8th percentile. Both Sweepfinder and H-scan detect elevated scores indicative of selection in the 100 kb region, though DARC is not an outlier (Table 1, S4-S6 Table). For example, using Sweepfinder, a method designed to detect recently completed hard selective sweeps based on the site frequency spectrum, the region is in the 97th percentile genome-wide in African populations. Similarly, using H-scan, a statistic designed to detect hard and soft sweeps via pairwise homozygosity tract lengths, we find the DARC region in the 95th percentile. We note however that accumulation of diversity and elevated recombination rate (average rate 3.33 cM/MB in 5kb region) may reduce the power of these statistics.

We also compared extended haplotype homozygosity (EHH) [50] and integrated haplotype score (iHH) [27] in the region for each of the three allelic classes (S1 Fig.). EHH patterns between populations of the same continent show strong similarity, consistent with the low levels of population structure observed within the continents. EHH in the FY*B samples decreases rapidly with genetic distance, while the FY*A and FY*O samples show higher levels linkage disequilibrium. When examining EHH separately for each of the two major FY*O haplotype backgrounds, we find increased linkage disequilibrium, as expected.

Finally, we screened ancient human genomes for the presence of the DARC alleles. We find no evidence for the FY*O mutation, consistent with the absence of genomes from sub-Saharan Africa in currently available ancient DNA datasets. The archaic hominin genomes of the Denisovan and Altai Neandertal carry the ancestral FY*B allele, while an ancient Ethiopian genome dated at 5,000 years old is a FY*A/FY*B heterozygote [51–53]. Additionally, we find that Ust’-Ishim, a 45,000 years old individual from Siberia [54] is also heterozygous for FY*A.

#### Evidence of positive selection at FY*A

Evidence for positive selection at the FY*A allele is currently under debate; binding assays show decreased binding of *P.vivax* to FY*A [13], though studies of the incidence of clinical malaria reach differing conclusions [12–14,16]. Despite this debate, it exhibits strong population differentiation and structure. FY*A is present at high frequency in Europe, Asia and southern Africa, but is conspicuously absent from the rest of sub-Saharan Africa. Similar to FY*O, FY*A has a very high *F*_*st*_ (99.99th percentile); however, selection scans based on the site frequency spectrum and linkage disequilibrium fail to detect selection (Table 1, S6 and S7 Tables). In Asian samples, which are about 90% FY*A, H-scan is in the 51st percentile, while Sweepfinder is slightly elevated to the 85th percentile.

We further analyzed the frequency trajectory of FY*A over time utilizing ancient genomes. We find that FY*A maintains a 30-50% frequency in our samples throughout most time periods and geographic regions, indicating that FY*A was already common in Eurasia as early as the Upper Paleolithic (S3 Fig). We note these frequencies are substantially lower than those observed in contemporary East Asian populations. However, most of the Bronze Age Asian samples are from the Altai region in Central Asia, which have been shown to derive a large fraction of their ancestry from West Eurasia sources [55]. We also note that the only published ancient African (Ethiopian) genome is heterozygous for the FY*A allele, indicating FY*A was likely not introduced into East Africa due to recent back migration [53].

### Inference of T_MRCA_ of FY*O and FY*A

We inferred the time to most recent common ancestor (T_MRCA_) of FY*A and FY*O based on the average number of pairwise differences between haplotypes. This method assumes a star-like phylogeny and no recombination [56]. Though FY*A and FY*O are approximately star-like (S2 Fig), allele age estimation is complicated by recombination within and between allelic classes. To address this caveat, we limited our calculations to the non-recombining region for each pair of haplotypes (see Methods). For FY*O, T_MRCA_ estimates were calculated separately for the two most common haplotypes.

We estimate the major FY*O haplotype class to be 42,183 years old (95% CI: 34,100 – 49,030) and the minor haplotype class to be 56,052 years old (95% CI: 38,927 – 75,073) (Table 2, S8-S10 Table). For the FY*A allele, the allele age was estimated as 57,184 years old (95% CI: 47,785 — 64,732). Variation between population-specific T_MRCA_ estimates was low. Additionally, we find that Ust’-Ishim, a 45,000 years old individual from Siberia [54] is heterozygous for FY*A. Under the assumption of no recurrent mutations, this would set a minimum age of 45,000 years for the FY*A mutation.

**Table 2.** T_MRCA_ results

|  |  | Estimate (years) | 95% CI (years) |
| --- | --- | --- | --- |
| FY*O | Major | 42,183 | 34,100 – 49,030 |
|  | Minor | 56,052 | 38,927 – 75,073 |
| FY*A |  | 57,184 | 47,785 – 64,732 |

T_MRCA_ results assume 25 year generation time and 1.2 * 10^−8^ mutations per base pair per generation. Confidence intervals are calculated from 1000 bootstrapped samples.

### Mode and magnitude of positive selection on FY*O

FY*O’s two divergent haplotypes indicate it may have reached fixation in Africa via selection on standing variation. To investigate this, we utilized an Approximate Bayesian Computation (ABC) approach to estimate the magnitude of FY*O’s allele frequency at selection onset, followed by the selection coefficient (*s*) of FY*O.

To infer the magnitude of FY*O’s allele frequency at selection onset, we compared the posterior probability of five models of initial frequency at selection onset (*de novo* mutation (1/2N), 0.1%, 1%, 10%, 25%), utilizing a Bayesian model selection approach in ABC [57–59]. Briefly, for each model we ran 100,000 simulations centered on an allele with selection coefficient drawn from the distribution 10^*U*(-3,-0.5)^ and recorded statistics summarizing the data. We assumed an additive selective model, as empirical studies predict heterozygotes have intermediate protection against *P. vivax* infection [11, 12] and a selection start time similar to the FY*O major haplotype’s T_MRCA_ (40 kya). We investigate our power to distinguish between the different models utilizing cross validation. We show that we have high power to distinguish between *de novo* and higher initial frequencies, though there is some overlap between adjacent models (S1 Appendix). Utilizing a multinomial logistic regression method, we observed strong support for the 0.1% initial frequency model and low support all other models (posterior probabilities: *de novo* 0.0002; 0.1% 0.9167; 1% 0.0827; 10% 0.0000; 25% 0.0004)(S1 Appendix). We conclude selection on FY*O occurred on standing variation with a very low (0.1%) allele frequency at selection onset.

We next sought to infer the strength of the selective pressure for FY*O. We estimated FY*O’s selection coefficient via ABC and local linear regression, assuming an allele frequency at selection onset of 0.1%. We find we have reasonable power to accurately infer *s* from these simulations; estimated and true selection coefficients have an *r*^2^ value of 0.85 with a slight bias of regression to the mean (S1 Appendix). We estimate the selection coefficient to 0.043 (95% CI: 0.011 – 0.18) (Fig 3). This selection coefficient is similar to other loci inferred to have undergone strong selection in the human genome, including skin pigmentation and other malaria resistance alleles [26, 60, 61].

**Fig 3.**
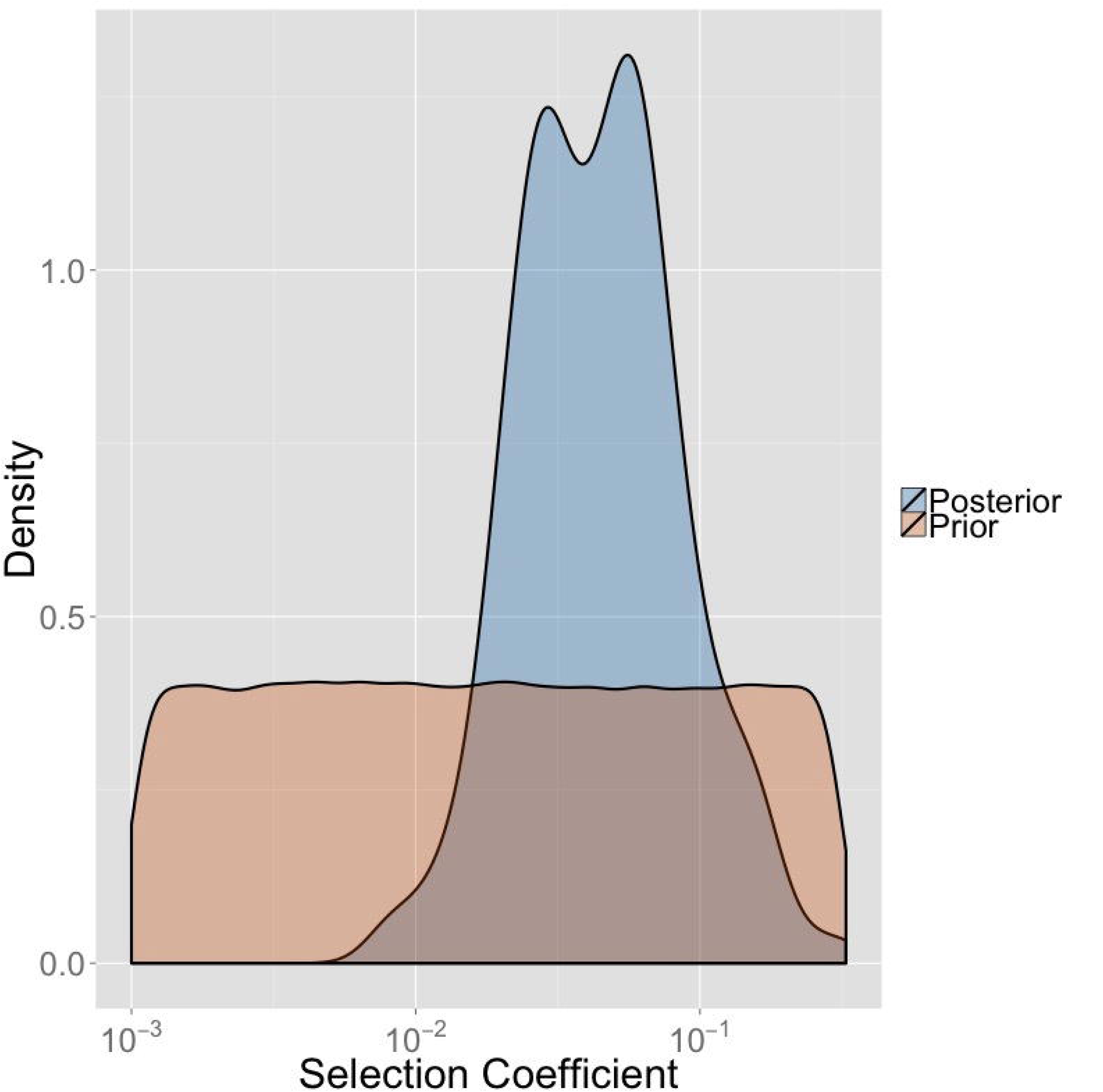
FY*O selection coefficient results Prior and posterior distributions of FY*O selection coefficient

To validate our model choice, we sampled selection coefficients from this posterior distribution and ran simulations with the initial frequency drawn from either 10^*U*(-5,-0.5)^ or *U*(0, 1). With the log-based prior distribution, we re-estimate the initial frequency at 0.15% (95% CI: 0.018 – 0.77%; S1 Appendix), closely fitting our inference. With the uniform prior distribution, we have much lower power to estimate initial allele frequency and we re-estimate the initial frequency at 6.86% (95% CI: −20.3 – 51.6%)(S1 Appendix). This is not surprising as it has previously been shown that it is very difficult to estimate initial frequency with this prior [62].

### Allelic Classes of Southern Africa

We also sought to understand the history of these alleles in southern Africa as, unlike equatorial Africa, malaria is not currently endemic in southwestern Africa and past climate was potentially unsuitable for malaria. Thus, we expect there was a lower or no selection pressure for FY*O or FY*A in this region. We analyzed sequences from the Bantu-speaking Zulu and indigenous ≠Khomani San. We find all three allelic classes are present in both populations (Zulu: FY*A 6%, FY*B 16%, FY*O 79%; ≠Khomani San: FY*A: 35%, FY*B 44%, FY*O 21%). The KhoeSan peoples are a highly diverse set of southern African populations that diverged from all other populations approximately 100 kya [63]. The Zulu population is a Bantu-speaking group from South Africa; southern Bantu-speakers derive 4 – 30% KhoeSan ancestry [64] from recent gene flow during the past 1,000 years. We first ask if the FY*O allele in the KhoeSan group represents recent gene flow from Bantu-speakers or whether FY*O has been segregating in southern Africa for thousands of years. We investigated global and local ancestry differences between FY*O carriers and non-carriers. We find a significant difference in genome-wide western African ancestry in ≠Khomani San FY*O carriers vs. non-carriers (17% average in FY*O carriers vs. 5.4% average in non-FY*O carriers, *p* = 0.014). We also find a significant enrichment of local Bantu-derived ancestry around the FY*O mutation in the ≠Khomani San FY*O carriers (*p* = 2.78*10^−12^; S4 and S5 Figs). Each of these factors indicate that FY*O was recently derived from gene flow into the ≠Khomani San population from either Bantu-speaking or eastern African groups. We then explored the relationship of FY*O in KhoeSan and Zulu samples to Bantu-speaking populations from equatorial Africa. A haplotype network of the ≠Khomani San FY*O carriers indicated that each 20kb haplotype was identical to a haplotype from populations further north (S6 Fig). We tested the Zulu FY*O samples as well, and found that they have identical, though more diverse, haplotypes than other Bantu-speaking populations (Fig 2). However, the increase in diversity may be due to calling biases and recombination between different allelic classes in the Zulus (see discussion).

We then sought to understand the prehistory of FY*A in southern Africa. The FY*A allele is common in San populations, despite its absence from equatorial Africa. We compared the FY*A haplotypes found in the ≠Khomani San and Zulu populations with FY*A haplotypes present in Asia and Europe to distinguish between three hypotheses. The FY*A mutation in southern Africa either was 1) segregating in the ancestral human population, 2) due to recent admixture from migrations ‘back to Africa’, or 3) arose convergently in a separate mutation event distinct from the European / Asian mutation. We find that Zulu FY*A haplotypes are highly diverse; some are identical to non-African FY*A haplotypes, while others are unique or ancestral (Fig. 2). Global ancestry results show no statistically significant difference between Bantu or KhoeSan ancestry in FY*A ≠Khomani San carriers and non-carriers (San: *p* = 0.85, Bantu: *p* = 0.101). Our local ancestry results indicate that FY*A carriers are significantly enriched for San ancestry around FY*A compared with non-carriers (*p* = 0.011). Our results support hypothesis (1), i.e. high ≠Khomani San FY*A haplotype diversity indicates FY*A has an ancient presence in southern Africa. Furthermore, as Bantu-speaking populations from equatorial Africa currently are exclusively FY*O, it is unlikely they transferred FY*A to KhoeSan after the Bantu expansion. Rather, the FY*A haplotypes in the Zulu are largely derived from admixture with the indigenous KhoeSan populations, or potentially very recent gene flow from European/Asian immigrants to South Africa.

### DARC in Great Ape Genomes

Great apes are known reservoirs of multiple species of *Plasmodium* [39]. DARC is an important infection factor not only for *P. vivax* but also for primate-related parasites such as *Plasmodium knowlesi* in macaques and *Hepatocystis kochi* in baboons [6,38,43]. Recent results suggest Japanese macaques are resistant to *P.vivax* potentially due to missense mutations in the binding region [65]. Furthermore, previous studies have found evidence for positive selection in DARC throughout the mammalian lineage based on sequence data from single individual great ape species [44]. We extend previous analyzes by including population level data for different great apes (chimpanzees, bonobos, and gorillas).

We extracted polymorphism data from the DARC gene region in chimpanzees, bonobos, and gorillas. All samples carry the ancestral FY*B allele. There were no mutations in the GATA-1 transcription box (the FY*O mutation location), and chimpanzees and bonobos exhibited no missense mutations in the *P. vivax* binding region of DARC (S11 Table). We also do not find the 100 kb region around the Duffy gene to be an outlier with respect to Tajima’s D, the number of pairwise differences, nor the number of segregating sites.

Notably, gorillas have one nonsynonymous SNP in the DARC *P.vivax* binding region, heterozygous in 3 of our 24 samples, which changes the same codon as the FY*A human mutation. The human FY*A mutation converts asparagine to glycine, while the gorilla mutation converts it to aspartic acid. However, asparagine is similar in charge and composition to aspartic acid, so functional impact of the mutation is unclear. Gorillas are also the only great ape species previously shown to have a fixed difference, also in the *P. vivax* binding region (V25A, [44]). This fixed difference has been shown to disrupt interaction between DARC and *P.vivax* ([66,67], though gorillas are still infected with *P.vivax*-like parasites [38]).

## Discussion

The FY*O allele in DARC is often cited as a quintessential example of positive selection in the human genome due to its biological implications and extreme continental *F*_*st*_. However, the population genetics and evolutionary history of this region remain understudied. Here, we infer the FY*O mutation in Africa to have undergone an ancient, soft selective sweep in equatorial Africa through multiple lines of evidence:

- Two divergent haplotypes forming separate star-like phylogenies
- Both divergent FY*O haplotypes found in hunter-gatherer and Bantu populations
- Low frequency of FY*O in southern Africa samples
- Ancient T_MRCA_ estimates of FY*O haplotypes
- Extreme population differentiation, but reduced signatures of selection in surrounding region
- ABC estimates of FY*O consistent with a low initial frequency and a high selection coefficient

In what follows, we explain how these different lines of evidence describe a complex picture for the evolution of this highly relevant locus for human evolution. First, we identify two divergent haplotypes carrying the FY*O mutation, an observation that is consistent with previous results [25,26]. These haplotypes, defined by four SNPs (one 600 bps upstream and three within 2500 bps downstream), are not compatible with a hard sweep model where one haplotype sweeps to fixation due to a positively selected *de novo* mutation. These two haplotypes both form star-like phylogenies and do not exhibit geographic structure in equatorial Africa, indicating both haplotypes were selected for in the same regions.

Second, identification of identical haplotypes in highly divergent African populations implies an ancient selective sweep before the complete divergence of these populations. The first line of evidence for this is that Baka and Mbuti populations have identical FY*O haplotypes in similar proportions as the Bantu populations. This is relevant because Baka and Mbuti are hunter-gatherer populations that diverged a long time ago from Bantu African populations (50 – 65 kya) as well as from each other (20 – 30 kya) [46,68–71]. Secondly, we observe low levels of admixture between these groups (Bantu admixture in Mbuti: 0 – 16%, Bantu admixture in Baka: 6.5 – 9.4%) [72,73]. However as many individuals were estimated to have no Bantu admixture, these identical haplotypes are unlikely to be due to recent gene flow. All together these observations are consistent with the mutation sweeping before or during the hunter-gatherer / Bantu split. This observation, along with the ancient T_MRCA_, is consistent with selection acting over this locus from ancient times, though even low levels of ancient gene flow may have resulted in its fixation due to its selection coefficient.

Third, FY*O’s much lower frequency in the ≠Khomani San, as well as other KhoeSan populations [9], indicates it may have had a lower selective pressure in southern Africa. This region’s past and current arid climate have made it a poor habitat for mosquitoes, reducing the associated risk of infection [74]. Furthermore, local and global ancestry results indicate FY*O may be due to recent gene flow into these populations, as ≠Khomani San FY*O carriers are significantly enriched for global Bantu ancestry and local Bantu ancestry in the FY*O region, relative to non-carriers.

Fourth, the confidence intervals of our FY*O T_MRCA_ estimates overlap the divergence times estimated for the hunter-gatherer / Bantu split, supporting the idea that the sweep occurred just before or during the split.

Fifth, the high *F*_*st*_, coupled with lower Sweepfinder and H-scan statistics, indicate an ancient sweep and/or selection on standing variation. A recent hard sweep in Africa would drastically reduce variation around the selected site (resulting in high homozygosity estimated from H-scan) and shift the site frequency spectrum to high and low frequency sites (inferred as selection by Sweepfinder). Instead, slightly lower H-scan and Sweepfinder statistics indicate more diversity and less extreme site frequency spectrum shifts than expected in a recent hard sweep. This may be due to an ancient sweep that had time to accumulate diversity and/or a sweep on standing variation that increased the frequencies of multiple diverse haplotypes. Selection on standing variation has been shown to have wider variance in relevant summary statistics and methods for detecting selection. The variance size depends on parameters such as allele frequency, time of selection, and strength of selection [75].

Sixth, ABC estimates initial FY*O frequency of magnitude 0.1% and selection coefficient 0.043 (95% CI:0.011 – 0.18). Though this initial frequency magnitude is very low, it drastically increases the probability that an allele of this selection coefficient will fix in the population, relative to a *de novo* mutation (see below).

### FY*O T_MRCA_ Estimates

We estimate T_MRCA_ of the most common haplotype class to be 42,183 years (95% CI: 34,100 – 49,030 years), after most estimates of the major predicted out-of-Africa expansion [76–79]. Previous estimates of the time of fixation of the FY*O mutation, based on lower density data, range from 9 – 63 kya (adjusted to our mutation rate and generation times) [26,80]. Other T_MRCA_ estimates ranging from 9 to 14 kya were calculated on microsatellites linked to FY*O [80], which seem to have underestimated the age of the mutation. Perhaps the most comprehensive work on this problem until now was by Hamblin and DiRienzo [26], who estimated the time to fixation of FY*O to be 63 kya (95% CI: 13,745 – 205,541 years; converted to our mutation rate). This is older than our estimates, but has overlapping confidence intervals. More recently, Hodgson et al. [61] estimated the time necessary for FY*O’s frequency to increase from 0.01 – 0.99 to be 41,150 years, based on an inferred selection coefficient in Madagascar.

The most common class of FY*O haplotypes (defined by four SNPs) exhibits a star-like genealogy (sign of exponential growth) and composes the vast majority of the haplotypes (86%). This leads us to consider this estimated T_MRCA_ to be the time of selection onset. If FY*O originated 42 kya, this would be consistent with selection occurring after the initial out-of-Africa expansion [76–78], explaining why FY*O is currently absent from European and Asian populations. For populations in Africa, the inferred absence of FY*O from KhoeSan populations (until the Bantu expansion) could be explained as the result of the KhoeSan having diverged from agriculturists and other hunter-gatherer populations around 100 – 150 kya [81]. Baka and Mbuti hunter-gatherers diverged from agriculturists 50 – 65 kya (though the confidence intervals of most estimates range from 20 – 120 kya) [46,68–71]. The time of divergence for these populations overlaps our confidence intervals of FY*O T_MRCA_ and selection onset. As we observe the same FY*O haplotypes throughout Africa, including in hunter-gatherer populations, selection may have occurred during the time these populations were diverging and still migrating between each other. In this scenario, it is still possible that even low gene flow may have led to FY*O fixation due to its selective advantage.

Our estimate of the minor FY*O haplotype class T_MRCA_, 56,052 years (95% CI: 38,927 – 75,073), has a higher variance and is older than the major FY*O haplotype class estimates. Larger confidence intervals are partially due to fewer minor haplotypes; 86% of FY*O haplotypes are in the major haplotype class. However, it is also possible that the ‘minor haplotype’ is actually composed of multiple low frequency haplotypes due to standing variation, which all increased in frequency during to selection. This minor haplotype class is also more ancestral that the major haplotype class, indicating the FY*O allele may have recombined onto the major haplotype class.

We find no ancient genomes with the FY*O allele, though this is not unexpected as there are no ancient African genomes currently available and the low observed frequencies of FY*O out of Africa are likely due to recent gene flow from North Africa (Tuscans in 1000G) [82,83]. Ancient DNA is subject to DNA damage, which enriches for mutations from guanine to adenine, indicating our estimates of FY*A frequency is likely an underestimate. We predict that as more ancient genomes are found in Africa, most of the FY*O mutations would be captured in the two major haplotypes.

### FY*A T_MRCA_ Estimates

We inferred FY*A to be an older mutation than FY*O, likely segregating throughout Africa before FY*O swept to fixation. We estimate FY*A to be 57,187 years old (95% CI: 47,785 – 64,732 years), 15,000 years older than the most common FY*O haplotype and overlapping estimates of the out-of-Africa expansion time [76–79]. Ancient DNA from a Paleolithic hunter-gatherer provides evidence that FY*A was already present in Eurasia by at least 45,000 years ago, thereby setting a lower bound for the age of the mutation. Its intermediate frequency in ≠Khomani San and Zulu populations, and similar haplotypic structure is consistent with FY*A existence in Africa at an appreciable frequency before the out-of-Africa expansion had occurred. The fact that southern Africa is at the opposite end of the expansion strongly supports this claim.

### Scaling Parameter Uncertainty

Our results are scaled with the mutation rate of 1.2 * 10^−8^ mutations / basepair / generation and a 25 year generation time. This mutation rate is supported by many previous whole-genome studies ([54,84–87]; range: 1 - 1.2 * 10^−8^ mutations / basepair / generation), but we are aware of recent studies suggesting a higher mutation rate that are either based on exome data ( [88–90]; range: 1.3 - 2.2 * 10^−8^ mutations / basepair / generation) or whole-genome data ( [91,92]; range: 1.61 - 1.66 * 10^−8^ mutations / basepair / generation). To take into account this uncertainty, we performed additional analyses using a mutation rate of 1.6 * 10^−8^ mutations / basepair / generation. With this higher rate, we estimate much later origins of the FY*O and FY*A mutations; specifically we would estimate the FY*O T_MRCA_ to be 32 kya (vs. 42 kya) and the FY*A T_MRCA_ to be about 43 kya (vs. 57 kya). It is important to consider that most quantities in population genetics are scaled by the mutation rate and effective population size. Therefore, any changes in the mutation rate result in changes not only in our T_MRCA_ estimates, but also in the timescale of the split between African and non-African populations. For example, a recent study of the divergence between African and non-Africans, estimates a median of divergence between 52-69 kya and a final split around 43 kya, using a mutation rate of 1.2 * 10^−8^ mutations / basepair / generation [78]. If we use a higher mutation rate of 1.6 * 10^−8^ mutations / basepair / generation the median divergence would be 39-52 kya with a final split around 33 kya. Thus, regardless of the mutation rate (and the corresponding demographic scenario), we estimate the FY*O mutation to have occurred soon after the estimated final split, while the FY*A mutation occurred when gene flow was still occurring.

### Data Processing

We note there is likely calling bias between the 1000 Genomes integrated dataset and the low coverage samples recalled in this paper. This is evidenced by multiple SNPs being present only in the recalled low-coverage data (Fig 2), despite some populations in the recalled data being highly diverged from each other but close to those in the 1000 genomes data. Due to this, most analyses were conducted by population. Our results show that this calling bias does not affect our conclusions. For example, T_MRCA_ estimates of 1000 Genomes populations and recalled samples are very similar [S8-S10 Tables].

### FY*O Initial Frequency and Selection Coefficient Estimations

FY*O’s two divergent, common haplotypes in Africa indicate it may have reached fixation due to selection on standing variation. We infer that the FY*O mutation underwent a selective sweep on standing variation with a selection coefficient comparable to some of the most strongly selected loci in the human genome [60]. Utilizing a Bayesian model selection approach implemented in an ABC framework, we find that FY*O likely rose to fixation via selection on standing variation; though the frequency of FY*O at selection onset was very low (0.1%). We estimate FY*O’s selection coefficient to be 0.043 (95% CI: 0.011 – 0.18), consistent with previous estimates (>0.002 in the Hausa [26], 0.066 in Madagascar [61]). The similarity of these results indicates FY*O may have a similar selective effect in diverse environments.

We have shown that there are multiple haplotypes (at least two main haplotypes) carrying the FY*O mutation in African populations that have diverged a long time ago, which is consistent with a scenario of selection on standing variation. Interestingly, we observe in our estimations that the most likely initial frequency for FY*O is only 0.1%. At first glance it would be reasonable to consider such a low initial frequency equivalent to a scenario of selection on a *de novo* mutation. In order to distinguish between the two possibilities we use the diffusion approximation by Kimura [93,94] to estimate the probability of fixation (equation 8 in [94]) and demonstrate that it is much more likely to reach fixation with an initial frequency of 0.1% than a scenario of a new mutation arising in the population. We find that an allele with selection coefficient 0.043 and initial frequency 0.001 has a 99.4% probability of fixing, while a *de novo* mutation with the same *s* has only an 8.2% probability of fixing. It is important to note that in our calculation the initial frequency (*p*) in the equation for the *de novo* mutation scenario is calculated using the effective population size, as opposed to the census population size. However, if we reasonably assume *N ≥ N_e_, p* is likely at least 0.1% in the population. This translates in our estimates for the probability of fixation of a *de novo* mutation being far more optimistic than expected if the ancestral African census population size was much larger than the effective size. This low initial frequency until 40 kya is consistent with FY*O’s absence from non-African present and ancient genomes.

### FY*O and FY*A Mutations and *P. vivax*

FY*O and FY*A are thought to be under positive selection due to *P. vivax*, a malaria-causing protozoan that infects red blood cells through the Duffy receptor. Individuals with the FY*O allele do not express the Duffy receptor in red blood cells resulting in immunity to *P.vivax* [6,10] and individuals with the FY*A allele may have lower infectivity rates [11–16]. Unlike *P. falciparum*, the most common and deadly malaria protozoan in Africa that uses multiple entry receptors, *P.vivax*’s one mode of entry allows the possibility of resistance with only one SNP.

Was *P.vivax* the selective pressure for the FY*O and FY*A mutations? *P. vivax* is currently prevalent in equatorial regions outside of Africa; however it is unknown if *P. vivax* has ever been endemic to Africa. There is an ongoing debate as to if *P.vivax* originated in Asia or Africa. Previously, it was thought *P.vivax* originated in Asia, as Asian *P.vivax* has the highest genetic diversity [40,95]. However, recent evidence shows global human-specific *P.vivax* forms a monophyletic cluster from *P. vivax-like* parasites infecting African great apes, suggesting an African origin [38].

Human-specific *P.vivax* sequences form a star-like phylogeny likely due to a relatively recent demographic expansion. Our T_MRCA_ estimates of human-specific *P.vivax* sequences are 70 – 250 kya (S12 Table), consistent with previous estimates (50 – 500 kya, [40,41,95]). The relative good overlap between T_MRCA_ of *P. vivax* and the T_MRCA_ of FY*O is consistent with the hypothesis of *P.vivax* being the selective agent responsible for the rise of FY*O in Africa. However, there are two possible scenarios that could explain the T_MRCA_ estimates for *P. vivax*. A first scenario is that the estimated T_MRCA_ of human *P.vivax* indicates the start of the association between host and parasite, thus marking the start of selective pressure on the host. A second scenario is that these estimates overlap the human out-of-Africa expansion times. It is possible that human-specific *P.vivax* expanded out of Africa with humans, resulting in the estimated T_MRCA_ for *P. vivax*. The human *P.vivax* currently in Africa could be from recent migration. Based on the phylogenies, it is unclear if human-specific African or Asian *P.vivax* are ancestral. Despite the recent observations of monophyletic relationships among all *P. vivax*, including African parasites, sufficient data remains that is inconsistent with *P. vivax* having an African origin. For example, the most closely related parasite to *P. vivax* is *P. cynomolgi*, a macaque parasite [38,40], and the most genetically diverse populations of *P. vivax* are in Asia and Melanesia [41,95]. Additionally, it is yet unclear if such a high selection coefficient is consistent with the fact that the general severity of *P. vivax* is currently much lower than that observed for *P. falciparum*, causing more morbidity than mortality. The combination of these observations lead us to suggest that further work is necessary to better understand the evolutionary history of *P. vivax* to reconcile the demographic scenarios that could have given rise to such a complex pattern.

All together, our results suggest that the evolutionary history of the FY*O mutation, a single SNP under strong selection in human populations, has been a complex one. Multiple haplotypes present in highly divergent African populations are consistent with selection on standing variation, shaping the evolution of this locus that was present in very low frequency in ancestral populations. Although more work needs to be done to understand how *P.vivax* may have shaped the evolution of this locus, our results provide a framework to evaluate the evolution of the parasite and formulate specific hypotheses for its evolutionary history in association with its human host.

## Materials and Methods

### Genetic Data and Processing

#### Modern Population Sequence Data

Data used in this study was retrieved from the African Genome Variation Project (AGVP, Zulu, Bagandans), Human Genome Diversity Project (HGDP, Mbuti [96]), the 1000 Genomes Project, as well as data sequenced in the lab (Sikora et al., In Prep) (Nzebi, Baka) and from [97] (≠Khomani San). Sequence data for 1000 genomes populations was retrieved from the phase 3 version 3 integrated phased call set (ftp://ftp-trace.ncbi.nih.gov/1000genomes/ftp/release/20130502/). Related individuals were removed (ftp://ftp-trace.ncbi.nih.gov/1000g/ftp/release/20130502/20140625 related individuals.txt).

SNPs from samples sequenced in-house, the AGVP and the HGDP were recalled together. Bam files from the AGVP were downloaded from the EGA archive via a data access agreement. Chromosome 1 bam files for all three data sources were cleaned with SamTools [98]. The following protocols were run to prepare the bam files: CleanSam.jar, FixMateInformation.jar, ValidateSamFile.jar, SortSam.jar, and MarkDuplicates.jar. We applied GATK [99] base quality score recalibration, indel realignment, and duplicate removal. We performed SNP discovery with GATK UnifiedGenotyper (default settings and min conf=10) and variant quality score recalibration according to GATK Best Practices and a tranche sensitivity threshold of 99% [100,101]. SNPs were phased and imputed by Beagle in two steps [102]. First, the 1000 Genomes sequences were used as a reference panel to phase and impute SNPs present in both datasets. Next, Beagle was run a second time without a reference panel to phase and impute remaining SNPs. The 20 kb region surrounding FY*O (chr1:159,164,683-159,184,683) was extracted from the 1000 Genomes data and merged with the recalled data. We identified 401 SNPs in the merged dataset. Analyses were restricted to biallelic SNPs. Derived and ancestral allelic states were determined via the human ancestor sequence provided by ensembl from the 6 primate EPO [103]. SNPs without a human ancestor were not included in analyses.

#### Ancient Genomes

Ancient genomes were processed as described in Allentoft et al. [55]. Briefly, we randomly sampled a high quality read for each ancient individual with coverage at the Duffy SNPs. Population allele frequencies were then estimated by combining multiple individuals into populations as in Allentoft et al. [55].

#### Great Ape Sequence Data

Great ape sequences mapped to their species-specific genomes from Prado-Martinez et al. [104] were utilized in this analysis. This included 24 chimpanzees (panTro-2.1.4), 13 bonobos (panTro-2.1.4), 24 gorillas (gorGor3), and 10 orangutans (ponAbe2). The DARC gene and 1kb surrounding region was extracted from each species based on Ensembl annotations: gorilla (chr1:138,515,328-138,517,811), and chimpanzees and bonobos (chr1:137,535,874-137,538,357) [105]. Orangutans were excluded from analyses because they have two regions orthologous to the human DARC gene (chr1:92,205,245-92,206,855, chr1 random:12,168,081-12,170,200,). SNP functionality was annotated by SNPEff [106].

### Population Structure Analyses

#### Haplotype Analyses

Median-joining networks were constructed via popArt [107].

#### Promoter Region Summary Statistics

Summary statistics (number of segregating sites, average number of pairwise difference, Tajima’s D) were calculated in the 750 bp promoter region upstream every genes in the 1000 Genomes integrated data via VCFtools [108]. The summary statistics from DARC’s promoter region were compare to all other promoter regions. Gene locations were extracted from ensembl release 72 [105].

#### Selection Summary Statistics

We analyzed methods in three main categories of selection detection: population differentiation (F_*ST*_), site frequency spectra (Sweepfinder [47,48]), and linkage disequilibrium (H-scan [49]). Genomic regions that have undergone a recent hard selective sweep are expected to have site frequency spectrums skewed toward rare and common variants, increased homozygosity and, if local adaptation, high population differentiation. Summary statistics were calculated for the fifteen 1000 Genomes populations.

Weir and Cockerham’s (1984) weighted *F*_*st*_ was calculated in VCFtools [108,109]. Sweepfinder, a method designed to detect recent hard selective sweeps based on the site frequency spectrum was ran via the SweeD software [47,48]. H-scan, a statistic designed to detect hard and soft sweeps [49], measures the average length of pairwise homozygosity tracts in a population. By utilizing pairwise homozygosity tracts, this method can detect soft sweeps, sweeps that have resulted in multiple haplotypes reaching high frequency. The default distance method was used (-d 0) and the maximum gap length between SNPs was set to 20kb. To calculate recombination adjusted results, recombination rates from deCODE [110] were lifted over from hg18 to hg19. We limited comparisons to regions with average recombination rates within 25% of the DARC region’s recombination rate. EHH was calculated via the R package rehh [111].

#### Inference of T_MRCA_

We estimated the T_MRCA_ of the FY*A and FY*O mutations through a method based on the average number of pairwise differences between two haplotypes [56]. We used the equation, 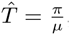, where *π* is the average number of pairwise differences per base pair in the sample and *μ* is the mutation rate per year per base pair. We assumed a mutation rate of 1.2 * 10^−8^ mutations per basepair per generation and a generation time of 25 years. Analyses were restricted to individuals homozygous for FY*B, FY*A or FY*O due to phase uncertainty. Regions were limited to “callable” sequence based on the 1000 genomes strict mask. 18,333 basepairs were callable in the 20kb region. Standard error estimates were calculated by 1000 bootstrap estimates with replacement.

This T_MRCA_ method calculates the average time to most recent common ancestor between two haplotypes in the sample. It assumes a star-like phylogeny and no recombination. Our phylogenies are close to star-like (S2 Fig) and Slatkin and Hudson [56] show that near star-like phylogenies, with *N*_0_ * *s* >> 0, result in valid allele age estimates. We estimate the T_MRCA_ of FY*O’s two major haplotypes separately as their deep divergence would strongly violate the star-like phylogeny requirement. We focused on a star-like phylogenetic method, as opposed to the coalescent, as the latter does not take into account selection, an apparently strongly influencing effect in this region, and thus would result in an artificially much older T_MRCA_ estimate.

We developed a variation of this method to account for recombination exhibited between allelic classes. For each pair of haplotypes, we identified the maximum region around the focal SNP with no signs of recombination between these haplotypes and haplotypes of other allelic classes. To identify this region, we expanded out from the focal SNP until we identified a SNP that was segregating both in the haplotype pair and in any samples in other allelic classes. This SNP is identified as a potential recombinant. The region for comparison is then set to the region between the two farthest nonrecombinant SNPs on each side plus half the region between the last nonrecombinant SNP and the first potential recombinant SNP. To calculate pairwise T_MRCA_, we count the number of nucleotide differences between the two haplotypes in this region. All pairwise T_MRCA_ estimates are then averaged to estimate sample T_MRCA_.

Minimum and maximum region sizes were also set. The minimum total sequence length was set to 3,000 basepairs, to ensure the expected number of mutations is at least one. This is important because if, for example, the SNPs adjacent to the focal SNP are both potential recombinants, the estimate allele age from these haplotypes would be 0, biasing the estimate to a more recent time. A maximum region size is set because the signature of selection decays as distance increases from the focal SNP, likely due to unseen recombination events. The maximum distance on each side was set to the distance in which EHH fell below 0.5 or 0.66 (FY*O 0.5: 3,322 bps upstream, 3,034 bps downstream; FY*O 0.66: 2,640 bps upstream, 3,034 bps downstream; FY*A 0.5 and 0.66: 4,358 bps upstream, 1,176 bps downstream). In most cases, small variations in the size of the selected region have little effect on the results; however, it did result in two very different estimates for the FY*O minor haplotype due to a common SNP included in the larger region size. The estimates with the EHH 0.66 cutoff is 56,052 years (95% CI: 38,927 – 75,073), while with the EHH 0.5 cutoff is 141,692 years (95% CI: 117,979 – 164,918).

#### FY*O Initial Frequency and Strength of Selection

To estimate FY*O’s selection coefficient and initial frequency at selection onset in equatorial Africa (based on the LWK population), we utilized an Approximate Bayesian Computation (ABC) approach in two steps: (1) we identified the best model of FY*O frequency at selection onset and (2) we estimated the selection coefficient assuming that initial frequency.

Inference was based on simulations, via *msms* [112], of 5 kb sequences centered on a selected allele with the African demographic model infered in Gravel et al. [76]. We assume a constant additive model of selection starting 40 kya (rounded T_MRCA_ of major FY*O haplotype class). For all simulations the prior distribution for the selection coefficient was *s* = 10^*U*(10^−0.5^-10^−3^)^. The recombination rate was inferred from the average for the 5kb region from the deCODE map (3.33 cM/MB) [110]. We assumed a mutation rate equal to 1.2 * 10-mutations per base pair per generation and a generation time of 25 years.

First, we utilized a Bayesian model selection approach in an ABC framework to estimate the magnitude of FY*O’s initial frequency at selection onset (implemented in the R package *abc* [57,113,114]). We compared five models of the initial FY*O frequency (*de novo* (1/2N), 0.1%, 1%, 10%, 25%). We ran 100,000 simulations for each model and recorded summary statistics: *π* (average number of pairwise differences), number of segregating sites, Tajima’s D, Fay and Wu’s *θ*_*H*_, number of unique haplotypes, linkage disequilibrium (average EHH centered on the selected site at the two ends of the sequences and iHH), allele frequency statistics (number of fixed sites, singletons, doubletons, singletons / fixed sites), H statistics [115] (H1, H2, H12, H2/H1), and final frequency of the selected allele. Summary statistics were centered, scaled, and transformed with PLS-DA to maximize differences between models, and we retained the top 5 PLS-DA components (mixOmics R package [116], similar to [62]). We then utilized a multinomial logistic regression method with a 1% acceptance rate to estimate the posterior probability of each model.

Second, based on the model with the highest posterior probability (initial frequency: 0.1%), we estimated the selection coefficient using ABC and local linear regression. We ran 200,000 simulations and utilized the most informative summary statistics, as determined via information gain: number of segregating sites, number of mutations with more than two copies, number of fixed sites, (number of singletons) / (number of fixed sites), H1, H2, H12, number unique haplotypes, average EHH at ends, and the final frequency of the selected allele. We centered, scaled, and transformed these statistics with PCA, retaining PCs that explained 95% of the variance. Last, we estimated the posterior distribution with a logistic regression model and a 1% acceptance rate.

#### Allelic Classes in Southern Africa

This analysis utilized data from the Zulu [64] (Omni 2.5 array and low coverage sequence data, re-called with the rest of the African samples) and ≠Khomani San (550K and Omni Express and Omni Express Plus) and exome data). Exome data along with SNP array data (550k, Omni Express and Omni Express Plus) were merged with the HGDP set for the network analysis. We examined 84 KhoeSan and 54 Human Genome Diversity Panel (HGDP) individuals from 7 different populations [97]. There were 8 Pathan, 8 Mbuti Pygmy, 8 Cambodian, 8 Mozabite, 8 Yakut, 8 Mayan and 6 San individuals in the HGDP data set. The HGDP genotype data used in this study was acquired from Dataset 2 Stanford University and contained about 660,918 tag SNPs from Illumina HuHap 650K [117]). Exome data of the HGDP data set was previously sequenced and used in our analysis. Single nucleotide polymorphism (SNP) array/genotype and exome data were merged using PLINK. The SNP array platforms were merged as follows: HGDP650K, KhoeSan 550K OmniExpress and OmniExpressExomePlus. All individuals in the data set had full exome data and SNPs with a missing genotype rate more than 36% were filtered out of the data set.

Global San ancestry percentages were calculated from array data via ADMIXTURE [118]. For the ≠Khomani San samples, Europeans and a panel of 10 African populations from each major geographic region were used as potential unsupervised source populations. As the array data did not include rs2814778 or rs12075, these alleles were acquired from the corresponding exome data for each individual. Zulu global ancestry percentages are from [64] and FY status was determined from the corresponding sequence data. Only samples with matching identification numbers for the array and sequence data were included.

Local ancestry was determined using RFMix v1.5.4 [119]. For the ≠Khomani San samples, input files were array specific phase files, Omni Express and 550k, with three potential ancestral populations: (LWK) Bantu-speaking Luhya from Kenya, (CEU) western European, and (SAN) Namibian San. For the Zulu samples, we first merged and phased Omni2.5 genotype data for the two reference populations (Luhya (LWK) from Kenya and Nama (Khoe) from southern Africa) and the admixed population (Zulu). The Luhya data was downloaded from 1000 Genomes Project phase 3 (100 individuals) and the Nama genotype data is in preparation [Liu et al., In Prep] (102 individuals). The Zulu Omni2.5 file was downloaded from the African Genome Diversity Project and contained 100 individuals. Files were merged with PLINK and sites with missing genotype rate greater than 10% were filtered out. SHAPEIT v2.r790 was used to phase this merged data set [120]. For further phasing accuracy, family information was included for the Nama individuals and the – duohmm option was used when running the phase command; there were 7 duos and 1 trio included in our data set. After phasing, related Nama individuals were removed and only the Nama individuals with limited admixture were kept as the San reference for input into RFMix. When running RFMix, the PopPhased option was selected in the command; this option re-phases the original data, correcting haplotype phasing. Additionally, the command was run with two iterations. Local ancestry around the coding region of Duffy was extracted and plotted. A similar procedure was used to call local ancestry for the ≠Khomani San population using RFMix v1.5.4 [96].

We also constructed a median joining network (using Network [121]), for the 20kb region centered on the FY*O mutation. Site-specific weights were determined based on GERP conservation score. GERP scores were obtained from the UCSC genome browser (http://hgdownload.cse.ucsc.edu/ gbdb/hg19/bbi/All hg19 RS.bw) based on an alignment of 35 mammals to human. The human hg19 sequence reference allele was not included in the calculation of GERP RS scores. SNPs with an extremely negative GERP score (-5 or lower) were down-weighted to 5, SNPs with a GERP score higher than 3 were up-weighted to 15, and SNPs with a GERP score in-between these values were weighted to 10. The FY*O mutation was given a weight of 10, though it had a GERP score of 4.27. Maximum parsimony was used post calculation to clean the network by switching off unnecessary median vectors. The resulting network was drawn and edited in DNA publisher [121].

#### T_MRCA_ of *P.vivax* genes

We estimated the T_MRCA_ of human-specific *P. vivax* gene sequences from Liu et al. [38]. We assumed a star-like phylogeny and used the same pairwise differences equation as in the FY*O/FY*A estimates to calculate the T_MRCA_ of each *P.vivax* gene. We assumed a mutation rate of 5.07 * 10^−9^ basepairs per generation and a generation time of 1 year [95].

## Acknowledgments

The authors thank Dmitri Petrov and Philipp Messer for their thoughtful discussion about summary statistics and H-scan.

**Supporting Information**

**S1 Appendix**

**Supporting methods**. Elaboration on the initial frequency & selection coefficient estimator.

**S1 Fig**

**EHH plots by allelic type** EHH plots for the 20kb region surrounding the FY*O mutation. A) FY*O samples centered on FY*O mutation B) FY*A samples centered on FY*A mutation C) FY*B samples centered on FY*O mutation D) FY*B samples centered on FY*A mutation

**S2 Fig**

**Genetree genealogies** Geneology from Genetree of the 5kb region around FY*O. Dots indicate mutations and bottom numbers indicate number of samples with that haplotype. A) Geneology of FY*A samples from CHB population. B) Geneology of FY*O samples from LWK population.

**S3 Fig**

**Allele frequencies over time and space** Paleo: paleolithic; Hunter: hunter-gatherer; neol: neolithic; baEur: Bronze Age Europe; baStep: Bronze Age Steppe region; baAsia: Bronze Age Asia; ir: Iron Age. Sequences from Allentoft et al. (2015)

**S4 Fig**

**Local ancestry around FY*O mutation in ≠Khomani San samples** A) Homozygous FY*B samples B) Homozygous FY*O samples C) Homozygous FY*A samples D) FY*O/FY*B samples E) FY*A/FY*B samples F) FY*A/FY*O samples

**S5 Fig**

**Local ancestry around FY*O mutation in Zulu samples** There were no homozygous FY*A samples. A) Homozygous FY*B samples B) Homozygous FY*O samples C) FY*B/FY*O samples D) FY*A/FY*O samples

**S6 Fig**

**Network image of 10 kb on either side of the Duffy locus**. Weights are based on GERP conservation score. Asterisk indicates the root of the network. Blue circles indicate FY*O haplotypes.

**S1 Table**

**Samples included in study**

**S2 Table**

**Allele frequencies by population**

**S3 Table**

**Nucleotide diversity statistics**. Nucleotide diversity statistics in the 5kb, 10kb, and 20kb region surrounding the FY*O mutation.

**S4 Table**

**Promoter region summary statistics**. Summary statistics were calculated in 750 bp region upstream from DARC and compared to the 750 bp region upstream from all other genes in genome in each population. Summary statistics calculated: number of segregating sites (s), number of pairwise differences (*π*), and Tajima’s D. We quantified the percentile in the genome (Per.), median, and 95% confidence interval (CI).

**S5 Table**

**F_*ST*_ statistics**. Weir and Cockerham’s weighted F_*ST*_ was calculated for each SNP in the genome and for 5 kb, 10 kb, and 20 kb windows. F_*ST*_ result and its percentile in the genome is reported for all fifteen 1000 Genomes populations.

**S6 Table**

**Sweepfinder statistics**. We report the likelihood that the Duffy region (20 kb and 100 kb) underwent a recent hard selective sweep. This likelihood is compared to likelihoods from all other regions in the genome, as well as regions with average recombination rates within 25% of the Duffy region’s recombination rate.

**S7 Table**

**H-scan statistics**. We report the maximum H-scan score for the Duffy region (20 kb and 100 kb). This score is then compared to the max score from all other regions in the genome, as well as regions with average recombination rates within 25% of the Duffy region’s recombination rate.

**S8 Table**

**T_MRCA_ results for FY*O major haplotype**. Results for the T_MRCA_ of FY*O major haplotype by population. Results assume 25 year generation time and mutation rate of 1.2 * 10^−8^ mutations per basepair per generation. Confidence intervals are calculated from 1000 bootstrapped samples.

**S9 Table**

**T_MRCA_ results for FY*O minor haplotype**. Results for the T_MRCA_ of FY*O minor haplotype by population. Results assume 25 year generation time and mutation rate of 1.2 * 10^−8^ mutations per basepair per generation. Confidence intervals are calculated from 1000 bootstrapped samples.

**S10 Table**

**T_MRCA_ results for FY*A haplotype**. Results for the T_MRCA_ of FY*A by population. Results assume 25 year generation time and mutation rate of 1.2 * 10^−8^ mutations per basepair per generation. Confidence intervals are calculated from 1000 bootstrapped samples.

**S11 Table**

**Great ape DARC nonsynonymous mutations**. All nonsynonymous mutations segregating in the DARC gene region in gorillas, chimpanzees, and bonobos.

**S12 Table**

**T_MRCA_ of *Plasmodium vivax* genes**

## REFERENCES

1. Haldane J. Disease and evolution. Ric Sci Suppl. 1949;19:68–76.

2. Kwiatkowski DP. How malaria has affected the human genome and what human genetics can teach us about malaria. Am J Hum Genet. 2005;77(2):171–192.

3. Gething PW, Elyazar IRF, Moyes CL, Smith DL, Battle KE, Guerra CA, et al. A long neglected world malaria map: Plasmodium vivax endemicity in 2010. PLoS Negl Trop Dis. 2012;6(9):e1814.

4. Howes RE, Patil AP, Piel FB, Nyangiri OA, Kabaria CW, Gething PW, et al. The global distribution of the Duffy blood group. Nat Commun. 2011;2:266.

5. Cutbush M, Mollison PL, Parkin DM. A new human blood group. Nature. 1950;165:188–189.

6. Miller LH, Mason SJ, Clyde DF, McGinniss MH. The resistance factor to Plasmodium vivax in blacks: the Duffy-blood-group genotype, FyFy. N Engl J Med. 1976;295(6):302–304.

7. Nurse G, Lane A, Jenkins T. Sero-genetic studies on the Dama of South West Africa. Ann Hum Biol. 1976;3(1):33–50.

8. Nurse GT, Jenkins T. Serogenetic studies on the Kavango peoples of South West Africa. Ann Hum Biol. 1977;4(5):465–478.

9. Nurse G, Botha M, Jenkins T. Sero-genetic studies on the San of south West Africa. Hum Hered. 1977;27(2):81–98.

10. Tournamille C, Colin Y, Cartron JP, Le Van Kim C. Disruption of a GATA motif in the Duffy gene promoter abolishes erythroid gene expression in Duffy–negative individuals. Nature Genet. 1995;10(2):224–228.

11. Kasehagen LJ, Mueller I, Kiniboro B, Bockarie MJ, Reeder JC, Kazura JW, et al. Reduced Plasmodium vivax erythrocyte infection in PNG Duffy-negative heterozygotes. PLoS One. 2007;2(3):e336.

12. Weber SS, Tadei WP, Martins AS. Polymorphism of the Duffy blood group system influences the susceptibility to Plasmodium vivax infection in the specific area from Brazilian Amazon. Brazilian Journal of Pharmacy. 2012;93(1):33–37.

13. King CL, Adams JH, Xianli J, Grimberg BT, McHenry AM, Greenberg LJ, et al. Fya/Fyb antigen polymorphism in human erythrocyte Duffy antigen affects susceptibility to Plasmodium vivax malaria. Proc Natl Acad Sci USA. 2011;108(50):20113–20118.

14. Chittoria A, Mohanty S, Jaiswal YK, Das A. Natural selection mediated association of the Duffy (FY) gene polymorphisms with Plasmodium vivax malaria in India. PloS one. 2012;7(9).

15. Carvalho TAA, Queiroz MG, Cardoso GL, Diniz IG, Silva ANLM, Pinto AYN, et al. Plasmodium vivax infection in Anajas, State of Para: no differential resistance profile among Duffy-negative and Duffy-positive individuals. Malar J. 2012;11:430.

16. Albuquerque SRL, Cavalcante FO, Sanguino EC, Tezza L, Chacon F, Castilho L, et al. FY polymorphisms and vivax malaria in inhabitants of Amazonas State, Brazil. Parasitol Res. 2010;106:1049–1053.

17. Woldearegai TG, Kremsner PG, Kun JFJ, Mordmüller B. Plasmodium vivax malaria in Duffy-negative individuals from Ethiopia. T Roy Soc Trop Med H. 2013;107(5):328–331.

18. Wurtz N, Mint Lekweiry K, Bogreau H, Pradines B, Rogier C, Ould Mohamed Salem Boukhary A, et al. Vivax malaria in Mauritania includes infection of a Duffy-negative individual. Malar J. 2011;10:336.

19. Cavasini CE, de Mattos LC, Couto AAD, Bonini-Domingos CR, Valencia SH, de Souza Neiras WC, et al. Plasmodium vivax infection among Duffy antigen-negative individuals from the Brazilian Amazon region: an exception? Trans R Soc Trop Med Hyg. 2007;101(10):1042–1044.

20. Ménard D, Barnadas C, Bouchier C, Henry-Halldin C, Gray LR, Ratsimbasoa A, et al. Plasmodium vivax clinical malaria is commonly observed in Duffy-negative Malagasy people. Proc Natl Acad Sci USA. 2010;107(13):5967–5971.

21. Ryan JR, Stoute JA, Amon J, Dunton RF, Mtalib R, Koros J, et al. Evidence for transmission of Plasmodium vivax among a duffy antigen negative population in Western Kenya. Am J Trop Med Hyg. 2006;75(4):575–581.

22. Sabeti P, Schaffner S, Fry B, Lohmueller J, Varilly P, Shamovsky O, et al. Positive natural selection in the human lineage. Science. 2006;312(5780):1614–1620.

23. Vallender EJ, Lahn BT. Positive selection on the human genome. Hum Mol Gen. 2004;13(suppl 2):R245–R254.

24. Wray GA. The evolutionary significance of cis-regulatory mutations. Nature Rev Genet. 2007;8(3):206–216.

25. Hamblin MT, Di Rienzo A. Detection of the signature of natural selection in humans: evidence from the Duffy blood group locus. Am J of Hum Genet. 2000;66(5):1669–1679.

26. Hamblin MT, Thompson EE, Di Rienzo A. Complex signatures of natural selection at the Duffy blood group locus. Am J of Hum Genet. 2002;70(2):369–383.

27. Voight BF, Kudaravalli S, Wen X, Pritchard JK. A map of recent positive selection in the human genome. PLoS Biol. 2006;4(3):446.

28. Sabeti PC, Varilly P, Fry B, Lohmueller J, Hostetter E, Cotsapas C, et al. Genome-wide detection and characterization of positive selection in human populations. Nature. 2007;449(7164):913–918.

29. Akey JM. Constructing genomic maps of positive selection in humans: Where do we go from here? Genome Res. 2009;19(5):711–722.

30. Zhou H, Hu S, Matveev R, Yu Q, Li J, Khaitovich P, et al. A Chronological Atlas of Natural Selection in the Human Genome during the Past Half-million Years. bioRxiv. 2015;p. 018929.

31. Williamson SH, Hubisz MJ, Clark AG, Payseur BA, Bustamante CD, Nielsen R. Localizing recent adaptive evolution in the human genome. PLoS Genet. 2007;3(6):e90.

32. Wang ET, Kodama G, Baldi P, Moyzis RK. Global landscape of recent inferred Darwinian selection for Homo sapiens. Proc Natl Acad Sci USA. 2006;103(1):135–140.

33. Tang K, Thornton KR, Stoneking M. A new approach for using genome scans to detect recent positive selection in the human genome. PLoS Biol. 2007;5(7):e171.

34. Kimura R, Fujimoto A, Tokunaga K, Ohashi J. A practical genome scan for population-specific strong selective sweeps that have reached fixation. PLoS one. 2007;2(3):e286–e286.

35. Kelley JL, Madeoy J, Calhoun JC, Swanson W, Akey JM. Genomic signatures of positive selection in humans and the limits of outlier approaches. Genome Res. 2006;16(8):980–989.

36. Shortt H, Garnham P, Malamos B. Pre-erythrocytic stage of mammalian malaria. Br Med J. 1948;1(4543):192.

37. Krief S, Escalante AA, Pacheco MA, Mugisha L, André C, Halbwax M, et al. On the diversity of malaria parasites in African apes and the origin of Plasmodium falciparum from Bonobos. PLoS Pathog. 2010;6(2):e1000765.

38. Liu W, Li Y, Shaw KS, Learn GH, Plenderleith LJ, Malenke JA, et al. African origin of the malaria parasite Plasmodium vivax. Nat Commun. 2014;5.

39. Prugnolle F, Durand P, Neel C, Ollomo B, Ayala FJ, Arnathau C, et al. African great apes are natural hosts of multiple related malaria species, including Plasmodium falciparum. Proc Natl Acad Sci USA. 2010;107(4):1458–1463.

40. Escalante AA, Cornejo OE, Freeland DE, Poe AC, Durrego E, Collins WE, et al. A monkey’s tale: the origin of Plasmodium vivax as a human malaria parasite. Proc Natl Acad Sci USA. 2005;102(6):1980–1985.

41. Cornejo OE, Escalante AA. The origin and age of Plasmodium vivax. Trends Parasitol. 2006;22(12):558–563.

42. Miller LH, Mason SJ, Dvorak JA, McGinniss MH, Rothman IK. Erythrocyte receptors for (Plasmodium knowlesi) malaria: Duffy blood group determinants. Science. 1975;189(4202):561–563.

43. Tung J, Primus A, Bouley AJ, Severson TF, Alberts SC, Wray GA. Evolution of a malaria resistance gene in wild primates. Nature. 2009;460(7253):388–391.

44. Demogines A, Truong KA, Sawyer SL. Species-specific features of DARC, the primate receptor for Plasmodium vivax and Plasmodium knowlesi. Mol Biol Evol. 2012;29(2):445–449.

45. Oliveira TYK, Harris EE, Meyer D, Jue CK, Silva Jr WA. Molecular evolution of a malaria resistance gene (DARC) in primates. Immunogenetics. 2012;64(7):497–505.

46. Patin E, Laval G, Barreiro LB, Salas A, Semino O, Santachiara-Benerecetti S, et al. Inferring the demographic history of African farmers and pygmy hunter-gatherers using a multilocus resequencing data set. PLoS Genet. 2009;5(4):e1000448.

47. Pavlidis P, Živković D, Stamatakis A, Alachiotis N. SweeD: likelihood-based detection of selective sweeps in thousands of genomes. Mol Biol Evol. 2013;30(9):2224–2234.

48. Nielsen R, Williamson S, Kim Y, Hubisz M, Clark A, Bustamante C. Genomic scans for selective sweeps using SNP data. Genome Res. 2005;15:1566–1575.

49. Schlamp F, van der Made J, Stambler R, Chesebrough L, Boyko AR, Messer PW. Evaluating the performance of selection scans to detect selective sweeps in domestic dogs. Mol Ecol. 2016;25(1):342–356.

50. Sabeti PC, Reich DE, Higgins JM, Levine HZP, Richter DJ, Schaffner SF, et al. Detecting recent positive selection in the human genome from haplotype structure. Nature. 2002;419(6909):832–837.

51. Meyer M, Kircher M, Gansauge M, Li H, Racimo F, Mallick S, et al. A high-coverage genome sequence from an archaic Denisovan individual. Science. 2012;338(6104):222–226.

52. Prüfer K, Racimo F, Patterson N, Jay F, Sankararaman S, Sawyer S, et al. The complete genome sequence of a Neanderthal from the Altai Mountains. Nature. 2014;505(7481):43–49.

53. Llorente MG, Jones E, Eriksson A, Siska V, Arthur K, Arthur J, et al. Ancient Ethiopian genome reveals extensive Eurasian admixture throughout the African continent. Science. 2015;p. aad2879.

54. Fu Q, Li H, Moorjani P, Jay F, Slepchenko SM, Bondarev AA, et al. Genome sequence of a 45,000-year-old modern human from western Siberia. Nature. 2014;514(7523):445–449.

55. Allentoft ME, Sikora M, Sjögren K, Rasmussen S, Rasmussen M, Stenderup J, et al. Population genomics of Bronze Age Eurasia. Nature. 2015;522(7555):167–172.

56. Slatkin M, Hudson RR. Pairwise comparisons of mitochondrial DNA sequences in stable and exponentially growing populations. Genetics. 1991;129(2):555–562.

57. Csillery K, Francois O, Blum MGB. abc: an R package for approximate Bayesian computation (ABC). Methods Ecol Evol. 2012;.

58. Beaumont MA, Zhang W, Balding DJ. Approximate Bayesian computation in population genetics. Genetics. 2002;162(4):2025–2035.

59. Pritchard JK, Seielstad MT, Perez-Lezaun A, Feldman MW. Population growth of human Y chromosomes: a study of Y chromosome microsatellites. Mol Biol Evol. 1999;16(12):1791–1798.

60. Hedrick PW. Population genetics of malaria resistance in humans. Heredity. 2011;107(4):283–304.

61. Hodgson JA, Pickrell JK, Pearson LN, Quillen EE, Prista A, Rocha J, et al. Natural selection for the Duffy-null allele in the recently admixed people of Madagascar. Proc R Soc B. 2014;281.

62. Peter BM, Huerta-Sanchez E, Nielsen R. Distinguishing between selective sweeps from standing variation and from a de novo mutation. PLOS Genet. 2012;p. e1003011.

63. Schlebusch CM, Skoglund P, Sjödin P, Gattepaille LM, Hernandez D, Jay F, et al. Genomic variation in seven Khoe-San groups reveals adaptation and complex African history. Science. 2012;338(6105):374–379.

64. Gurdasani D, Carstensen T, Tekola-Ayele F, Pagani L, Tachmazidou I, Hatzikotoulas K, et al. The African Genome Variation Project shapes medical genetics in Africa. Nature. 2015;517(7534):327–332.

65. Tachibana S, Kawai S, Katakai Y, Takahashi H, Nakade T, Yasutomi Y, et al. Contrasting infection susceptibility of the Japanese macaques and cynomolgus macaques to closely related malaria parasites, Plasmodium vivax and Plasmodium cynomolgi. Parasitol Int. 2015;64(3):274–281.

66. Tournamille C, Filipe A, Wasniowska K, Gane P, Lisowska E, Cartron J, et al. Structure–function analysis of the extracellular domains of the Duffy antigen/receptor for chemokines: characterization of antibody and chemokine binding sites. Brit J Haematol. 2003;122(6):1014–1023.

67. Tournamille C, Filipe A, Badaut C, Riottot M, Longacre S, Cartron J, et al. Fine mapping of the Duffy antigen binding site for the Plasmodium vivax Duffy-binding protein. Mol Biochem Parasitol. 2005;144(1):100–103.

68. Batini C, Lopes J, Behar DM, Calafell F, Jorde LB, Van der Veen L, et al. Insights into the demographic history of African Pygmies from complete mitochondrial genomes. Mol Biology Evol. 2011;28(2):1099–1110.

69. Veeramah KR, Wegmann D, Woerner A, Mendez FL, Watkins JC, Destro-Bisol G, et al. An early divergence of KhoeSan ancestors from those of other modern humans is supported by an ABC-based analysis of autosomal resequencing data. Mol Biol Evol. 2012;29(2):617–630.

70. Verdu P, Austerlitz F, Estoup A, Vitalis R, Georges M, Théry S, et al. Origins and genetic diversity of pygmy hunter-gatherers from Western Central Africa. Curr Biol. 2009;19(4):312–318.

71. Quintana-Murci L, Quach H, Harmant C, Luca F, Massonnet B, Patin E, et al. Maternal traces of deep common ancestry and asymmetric gene flow between Pygmy hunter–gatherers and Bantu-speaking farmers. Proc Natl Acad Sci USA. 2008;105(5):1596–1601.

72. Patin E, Siddle KJ, Laval G, Quach H, Harmant C, Becker N, et al. The impact of agricultural emergence on the genetic history of African rainforest hunter-gatherers and agriculturalists. Nat Commun. 2014;5.

73. Loh PR, Lipson M, Patterson N, Moorjani P, Pickrell JK, Reich D, et al. Inferring admixture histories of human populations using linkage disequilibrium. Genetics. 2013;193(4):1233–1254.

74. Gething PW, Van Boeckel TP, Smith DL, Guerra CA, Patil AP, Snow RW, et al. Modelling the global constraints of temperature on transmission of Plasmodium falciparum and P. vivax. Parasit Vectors. 2011;4(92):4.

75. Prezeworski M, Coop G, Wall JD. The signature of positive selection on standing genetic variation. Evolution. 2005;59(11):2312–2323.

76. Gravel S, Henn BM, Gutenkunst RN, Indap AR, Marth GT, Clark AG, et al. Demographic history and rare allele sharing among human populations. Proc Natl Acad Sci USA. 2011;108(29):11983–11988.

77. Schiffels S, Durbin R. Inferring human population size and separation history from multiple genome sequences. Nature genetics. 2014;.

78. Li H, Durbin R. Inference of human population history from individual whole-genome sequences. Nature. 2011;475(7357):493–496.

79. McEvoy BP, Powell JE, Goddard ME, Visscher PM. Human population dispersal “Out of Africa” estimated from linkage disequilibrium and allele frequencies of SNPs. Genome research. 2011;21(6):821–829.

80. Seixas S, Ferrand N, Rocha J. Microsatellite variation and evolution of the human Duffy blood group polymorphism. Mol Biol Evol. 2002;19(10):1802–1806.

81. Gronau I, Hubisz MJ, Gulko B, Danko CG, Siepel A. Bayesian inference of ancient human demography from individual genome sequences. Nature Genet. 2011;43(10):1031–1034.

82. Moorjani P, Patterson N, Hirschhorn JN, Keinan A, Hao L, Atzmon G, et al. The history of African gene flow into Southern Europeans, Levantines, and Jews. PLoS Genet. 2011;7(4):e1001373.

83. Botigué LR, Henn BM, Gravel S, Maples BK, Gignoux CR, Corona E, et al. Gene flow from North Africa contributes to differential human genetic diversity in southern Europe. Proceedings of the National Academy of Sciences. 2013;110(29):11791–11796.

84. Kong A, Frigge ML, Masson G, Besenbacher S, Sulem P, Magnusson G, et al. Rate of de novo mutations and the importance of father/’s age to disease risk. Nature. 2012;488(7412):471–475.

85. Campbell CD, Chong JX, Malig M, Ko A, Dumont BL, Han L, et al. Estimating the human mutation rate using autozygosity in a founder population. Nature genetics. 2012;44(11):1277–1281.

86. Ségurel L, Wyman MJ, Przeworski M. Determinants of mutation rate variation in the human germline. Annual review of genomics and human genetics. 2014;15:47–70.

87. Michaelson JJ, Shi Y, Gujral M, Zheng H, Malhotra D, Jin X, et al. Whole-genome sequencing in autism identifies hot spots for de novo germline mutation. Cell. 2012;151(7):1431–1442.

88. O’Roak BJ, Vives L, Girirajan S, Karakoc E, Krumm N, Coe BP, et al. Sporadic autism exomes reveal a highly interconnected protein network of de novo mutations. Nature. 2012;485(7397):246–250.

89. Neale BM, Kou Y, Liu L, Ma’Ayan A, Samocha KE, Sabo A, et al. Patterns and rates of exonic de novo mutations in autism spectrum disorders. Nature. 2012;485(7397):242–245.

90. Sanders SJ, Murtha MT, Gupta AR, Murdoch JD, Raubeson MJ, Willsey AJ, et al. De novo mutations revealed by whole-exome sequencing are strongly associated with autism. Nature. 2012;485(7397):237–241.

91. Lipson M, Loh P, Sankararaman S, Patterson N, Berger B, Reich D. Calibrating the Human Mutation Rate via Ancestral Recombination Density in Diploid Genomes. PLoS Genet;11(11):e1005550.

92. Palamara PF, Francioli LC, Wilton PR, Genovese G, Gusev A, Finucane HK, et al. Leveraging Distant Relatedness to Quantify Human Mutation and Gene-Conversion Rates. The American Journal of Human Genetics. 2015;97(6):775–789.

93. Kimura M. Some problems of stochastic processes in genetics. Ann of Math Stat. 1957;p. 882–901.

94. Kimura M. On the probability of fixation of mutant genes in a population. Genetics. 1962;47(6):713.

95. Mu J, Joy DA, Duan J, Huang Y, Carlton J, Walker J, et al. Host switch leads to emergence of Plasmodium vivax malaria in humans. Mol Biol Evol. 2005;22(8):1686–1693.

96. Uren C, Kim M, Martin AR, Bobo D, Gignoux CR, van Helden PD, et al. Fine-scale human population structure in southern Africa reflects ecological boundaries. bioRxiv. 2016;Available from: http://biorxiv.org/content/early/2016/02/03/038729.

97. Henn BM, Botigué LR, Peischl S, Dupanloup I, Lipatov M, Maples BK, et al. Distance from sub-Saharan Africa predicts mutational load in diverse human genomes. Proc Natl Acad Sci. 2105;(doi:10.1073/pnas.1510805112).

98. Li H, Handsaker B, Wysoker A, Fennell T, Ruan J, Homer N, et al. The sequence alignment/map format and SAMtools. Bioinformatics. 2009;25(16):2078–2079.

99. McKenna A, Hanna M, Banks E, Sivachenko A, Cibulskis K, Kernytsky A, et al. The Genome Analysis Toolkit: a MapReduce framework for analyzing next-generation DNA sequencing data. Genome Res. 2010;20(9):1297–1303.

100. DePristo MA, Banks E, Poplin R, Garimella KV, Maguire JR, Hartl C, et al. A framework for variation discovery and genotyping using next-generation DNA sequencing data. Nature Genet. 2011;43(5):491–498.

101. Auwera GA, Carneiro MO, Hartl C, Poplin R, del Angel G, Levy-Moonshine A, et al. From FastQ data to high-confidence variant calls: the genome analysis toolkit best practices pipeline. Curr Protoc Bioinformatics. 2013;p. 11–10.

102. Browning SR, Browning BL. Rapid and accurate haplotype phasing and missing-data inference for whole-genome association studies by use of localized haplotype clustering. Am J Hum Genet. 2007;81(5):1084–1097.

103. Cunningham F, Amode MR, Barrell D, Beal K, Billis K, Brent S, et al. Ensembl 2015. Nucleic Acids Res. 2015;43(D1):D662–D669.

104. Prado-Martinez J, Sudmant PH, Kidd JM, Li H, Kelley JL, Lorente-Galdos B, et al. Great ape genetic diversity and population history. Nature. 2013;499(7459):471–475.

105. Flicek P, Amode MR, Barrell D, Beal K, Billis K, Brent S, et al. Ensembl 2014. Nucleic Acids Res. 2013;p. gkt1196.

106. Cingolani P, Platts A, Wang LL, Coon M, Nguyen T, Wang L, et al. A program for annotating and predicting the effects of single nucleotide polymorphisms, SnpEff: SNPs in the genome of Drosophila melanogaster strain w1118; iso-2; iso-3. Fly. 2012;6(2):80–92.

107. popArt;. http://popart.otago.ac.nz.

108. Danecek P, Auton A, Abecasis G, Albers CA, Banks E, DePristo MA, et al. The variant call format and VCFtools. Bioinformatics. 2011;27(15):2156–2158.

109. Weir BS, Cockerham CC. Estimating F-statistics for the analysis of population structure. Evolution. 1984;p. 1358–1370.

110. Kong A, Thorleifsson G, Gudbjartsson DF, Masson G, Sigurdsson A, Jonasdottir A, et al. Fine-scale recombination rate differences between sexes, populations and individuals. Nature. 2010;467(7319):1099–1103.

111. Gautier M, Vitalis R. rehh: an R package to detect footprints of selection in genome-wide SNP data from haplotype structure. Bioinformatics. 2012;28(8):1176–1177.

112. Ewing G, Hermisson J. MSMS: a coalescent simulation program including recombination, demographic structure and selection at a single locus. Bioinformatics. 2010;26(16):2064–2065.

113. Beaumont MA. Joint determination of topology, divergence time, and immigration in population trees. Renfrew C Matsumura S, Forster P, editor, Simulation, Genetics and Human Prehistory, McDonald Institute Monographs. 2008;p. 134–1541.

114. Fagundes NJR, Ray N, Beaumont M, Neuenschwander S, Salzano FM, Bonatto SL, et al. Statistical evaluation of alternative models of human evolution. Proc Natl Acad Sci USA. 2007;104(45):17614–17619.

115. Garud NR, Messer PW, Buzbas EO, Petrov DA. Recent selective sweeps in North American Drosophila melanogaster show signatures of soft sweeps. PLoS Genet. 2015;11(2):e1005004.

116. Cao KL, Gonzalez I, Dejean S. mixOmics: Omics Data Integration Project; 2015. R package version 5.0-4. Available from: http://CRAN.R-project.org/package=mixOmics.

117. Li JZ, Absher DM, Tang H, Southwick AM, Casto AM, Ramachandran S, et al. Worldwide human relationships inferred from genome-wide patterns of variation. science. 2008;319(5866):1100–1104.

118. Alexander DH, Novembre J, Lange K. Fast model-based estimation of ancestry in unrelated individuals. Genome research. 2009;19(9):1655–1664.

119. Maples BK, Gravel S, Kenny EE, Bustamante CD. RFMix: a discriminative modeling approach for rapid and robust local-ancestry inference. Am J Hum Genet. 2013;93(2):278–288.

120. O’Connell J, Gurdasani D, Delaneau O, Pirastu N, Ulivi S, Cocca M, et al. A general approach for haplotype phasing across the full spectrum of relatedness. PLoS Genet. 2014;10(4):e1004234.

121. LTD FT. Network Publisher ver 2.0.0.1; 2013.

